# Color Data v2: a user-friendly, open-access database with hereditary cancer and hereditary cardiovascular conditions datasets

**DOI:** 10.1101/2020.01.15.907212

**Authors:** Mark J. Berger, Hannah E. Williams, Ryan Barrett, Anjali D. Zimmer, Wendy McKennon, Huy Hong, Jeremy Ginsberg, Alicia Y. Zhou, Cynthia L. Neben

## Abstract

Publicly-available genetic databases promote data sharing and fuel scientific discoveries for the prevention, treatment, and management of disease. In 2018, we built Color Data, a user-friendly, open access database containing genotypic and self-reported phenotypic information from 50,000 individuals who were sequenced for 30 genes associated with hereditary cancer. In a continued effort to promote access to these types of data, we launched Color Data v2, an updated version of the Color Data database. This new release includes additional clinical genetic testing results from more than 18,000 individuals who were sequenced for 30 genes associated with hereditary cardiovascular conditions, as well as polygenic risk scores for breast cancer, coronary artery disease, and atrial fibrillation. In addition, we used self-reported phenotypic information to implement the following four clinical risk models: Gail Model for five-year risk of breast cancer, Claus Model for lifetime risk of breast cancer, simple office-based Framingham Coronary Heart Disease Risk Score for ten-year risk of coronary heart disease, and CHARGE-AF simple score for five-year risk of atrial fibrillation. These new features and capabilities are highlighted through two sample queries in the database. We hope that the broad dissemination of this data will help researchers continue to explore genotype-phenotype correlations and identify novel variants for functional analysis, enabling scientific discoveries in the field of population genomics.

Database URL: https://data.color.com/

## INTRODUCTION

The use of next generation sequencing (NGS) technologies in research and clinical laboratories has led to a rapid increase of genetic data. However, there is a lack of publicly-available genetic data, especially paired genotypic-phenotypic data. In 2018, we launched Color Data, an open-access, cloud-based database containing genotypic and self-reported phenotypic information from 50,000 individuals who were sequenced for 30 genes associated with hereditary cancer (1). Color Data has already made an impact on the scientific community, being utilized in peer-reviewed publications (2,3) and presented as a resource to educators (4). Its user-friendly interface enables researchers to easily execute their own queries with self-serve filters and displays the results as text, tables, and graphs. Results can also be downloaded in different files formats for further analyses and shared via email or social media.

Another important consideration when designing and implementing the database was scalability for volume and integration of different data points. For the second release of Color Data (Color Data v2), we added new features to the existing hereditary cancer dataset as well as a new dataset of genotypic and self-reported phenotypic information related to hereditary cardiovascular conditions from more than 18,000 individuals. Importantly, the hereditary cardiovascular conditions dataset retains the same user-friendly interface, making it easy for researchers and scientists to explore a new disease area.

Here we describe updates made to the database, including changes to the cohort and the addition of new query filters and results such as family health history. We also added clinical risk models to Color Data v2, such as the Gail Model (5) and the Claus Model (6) for breast cancer, simple office-based Framingham Coronary Heart Disease Risk Score (7) for coronary heart disease, and CHARGE-AF simple score (8) for atrial fibrillation. These risk models are commonly used by healthcare providers in the clinic and are important tools to estimate risk. Recent work has demonstrated that polygenic scores can also accurately predict and stratify risk for common, complex diseases and can identify individuals who have magnitude of risk for disease similar to those with pathogenic or likely pathogenic variants (9,10). As such, Color Data v2 also includes polygenic scores for breast cancer, coronary artery disease, and atrial fibrillation (AF). We highlight the addition of the hereditary cardiovascular conditions dataset, clinical risk models, and polygenic scores through two sample queries in the database. To our knowledge, Color Data is the first database to include pre-calculated scores from clinical risk models and for polygenic risk, which can help researchers investigate the relationship between different types of risk factors, both genetic and non-genetic, for disease.

## MATERIALS AND METHODS

Design and implementation of the database were previously described in the flagship publication by Barrett, Neben et al. (1). All individuals included in Color Data v2 received a multi-gene NGS panel test from Color Genomics, Inc. (‘Color’, Burlingame, CA) for 30 genes associated with hereditary cancer. In addition, a subset of individuals also received multi-gene NGS panel testing for 30 genes associated with hereditary cardiovascular conditions. All individuals consented to have their genetic and self-reported phenotypic information appear in Color’s research database.

Laboratory procedures, bioinformatics analysis, and variant interpretation for the multi-gene panel tests were performed at Color (Burlingame, CA) under Clinical Laboratory Improvements Amendments (#05D2081492) and College of American Pathologists (#8975161) compliance as previously described (11). Bioinformatics analysis included the previously described 30 genes associated with hereditary cancer and was updated to include an additional 30 genes associated with hereditary cardiovascular conditions: *ACTA2, ACTC1, APOB, COL3A1, DSC2, DSG2, DSP, FBN1, GLA, KCNH2, KCNQ1, LDLR, LMNA, MYBPC3, MYH7, MYH11, MYL2, MYL3, PCSK9, PKP2, PRKAG2, RYR2, SCN5A, SMAD3, TGFBR1, TGFBR2, TMEM43, TNNI3, TNNT2*, and *TPM1*. Analysis, variant calling, and reporting focused on the complete coding sequence and adjacent intronic sequence of the primary transcript(s), unless otherwise indicated. In *APOB*, exon 1 was not analyzed, and variants of uncertain significance (VUS) were not reported. In *MYH7*, VUS were not reported for exon 27. In several genes, certain exons were not analyzed: exons 4 and 14 of *KCNH2*, exon 1 of *KCNQ1*, exon 11 of *MYBPC3*, exon 5 of *PRKAG2*, and exon 1 of *TGFBR1*.

Laboratory procedures and imputation for low coverage whole genome sequencing were performed at Color as previously described (12). Data from low coverage whole genome sequencing were used to calculate previously published polygenic scores for breast cancer (10), coronary artery disease (9), and atrial fibrillation (9). Each polygenic score was normalized using principal components analysis to account for the effects of population stratification. While polygenic scores have the highest performance in people of European ancestry, recent studies have demonstrated that they have stratification ability across global populations as well (13,14). To note, if users would like to view polygenic risk score results for a given query, they must select ‘Calculated’ in the polygenic risk score filter because only a subset of the individuals in the database have a calculated polygenic risk score. Individuals who do not have polygenic risk scores calculated are captured under the filter value ‘Unknown’. Other self-reported phenotypic and genotypic information from ‘Calculated’ and ‘Unknown’ individuals is included in other query results by default, unless otherwise selected.

Genotypic and self-reported phenotypic information were used in the following clinical risk models: Gail Model for five-year risk of breast cancer (5), Claus Model for lifetime risk of breast cancer (6), simple office-based Framingham Coronary Heart Disease Risk Score for ten-year risk of coronary heart disease (7), and CHARGE-AF simple score for five-year risk of atrial fibrillation (8). To note, only a subset of individuals have a risk score calculated. Individuals who do not have a risk score calculated are labeled as ‘Unknown’ if not enough information was provided to calculate a risk score or ‘Ineligible’ if they did not meet the model criteria. The eligibility criteria for risk models are as follows:

- Gail Model: female, age 35 - 85 years, no personal history of breast cancer, no likely pathogenic or pathogenic variants in a gene associated with hereditary breast cancer (*BRCA1, BRCA2, TP53, PTEN, STK11, CDH1, PALB2, CHEK2, ATM, NBN, BARD1*, and *BRIP1*)
- Claus Model: female, age 30 - 79 years, no personal history of breast cancer, no likely pathogenic or pathogenic variants in a gene associated with hereditary breast cancer (*BRCA1, BRCA2, TP53, PTEN, STK11, CDH1, PALB2, CHEK2, ATM, NBN, BARD1*, and *BRIP1*)
- Simple office-based Framingham Coronary Heart Disease Risk Score: age 30 - 74 years
- CHARGE-AF simple score: age 46 - 90 years

Risk score filter values for hereditary cancer are defined in Table 1 and for hereditary cardiovascular conditions in Table 2.

**Table 1.** New filter categories and filter values on the hereditary cancer dashboard.

| Filter categories | Filter values |
| --- | --- |
| Gail Risk Score | Elevated risk (age 35 - 49 years and risk $\geq 1.67\%$; age 50 - 59 years and risk $\geq 2.0\%$; age 60 - 69 years and risk $\geq 3.0\%$; age 70 - 74 years and risk $\geq 4.0\%$), Ineligible†, Non-elevated risk, Unknown* |
| Claus Risk Score | Elevated ($\geq 20\%$), Ineligible†, Non-elevated ($< 20\%$), Unknown* |
| BC Polygenic Risk Score‡ | Calculated, Unknown* |

BC, breast cancer. †Ineligible includes females who did not meet the model criteria and males. \*Unknown includes females who did not provide enough information to calculate risk scores. ‡Males included. Only displays normalized risk scores between -3 and 3.

**Table 2.**
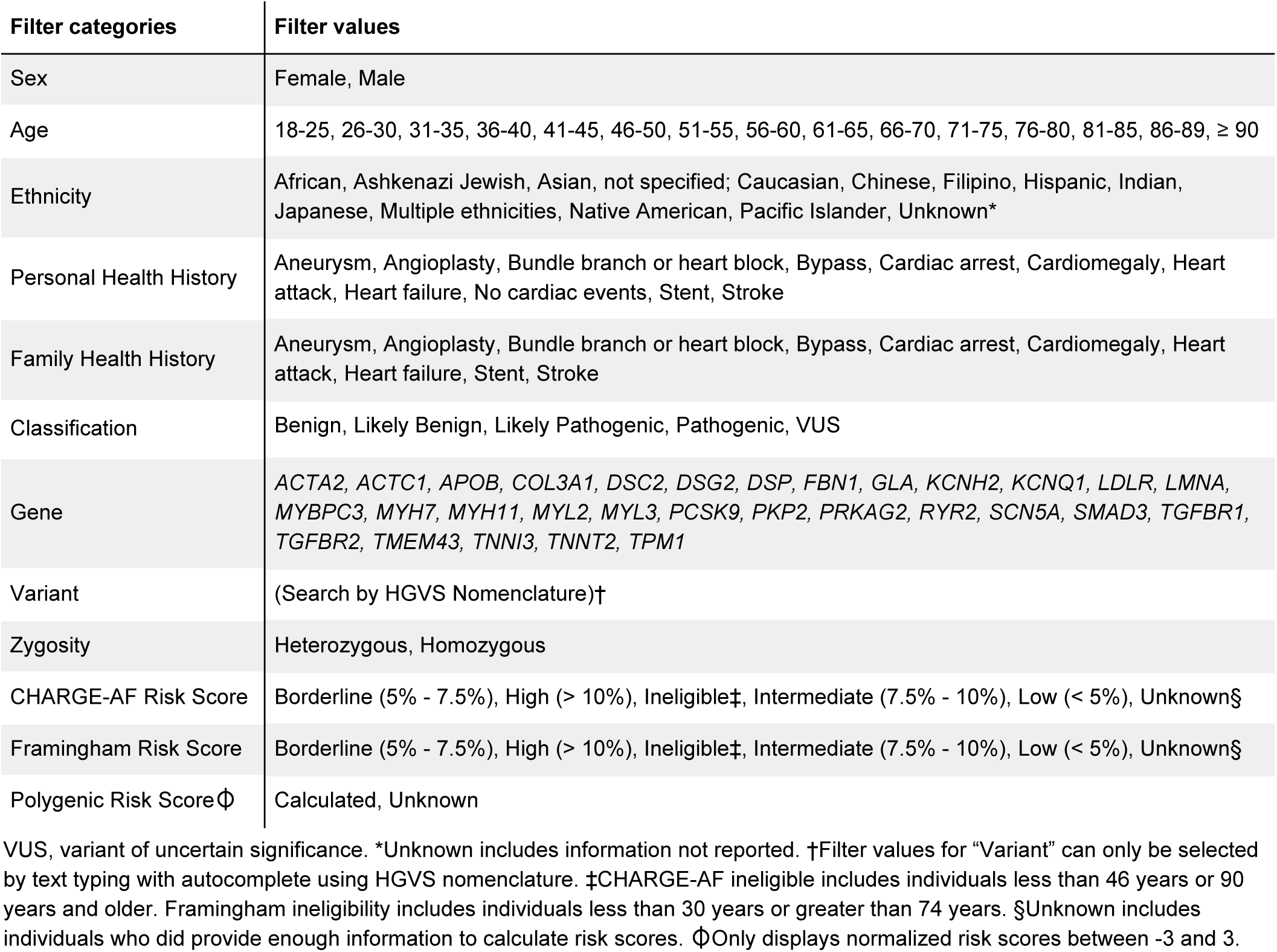
Filter categories and filter values on hereditary cardiovascular conditions dashboard.

## RESULTS

### Web interface

The Color Data home page (https://data.color.com/) links to two new query/result pages (hereafter referred to as “dashboards”): one for hereditary cancer and one for hereditary cardiovascular conditions. The links to three sample queries on the home page have been updated to demonstrate to users potential use cases of these dashboards as well as new query filters and results.

On the hereditary cancer dashboard (https://data.color.com/v2/cancer.html), users can apply the new query filters for family health history, risk models, and a polygenic risk score. These new filter categories and filter values are listed in Table 1. To note, the ‘AND’ logic for filter categories and ‘OR’ logic for filter values within categories still apply. New query results for hereditary cancer include ‘Gail Risk Score - 5 Year Risk of Breast Cancer’, ‘Claus Risk Score - Lifetime Risk of Breast Cancer’, and ‘Breast Cancer (BC) Polygenic Risk Score’. For ‘Breast Cancer (BC) Polygenic Risk Score’, results will only be displayed if a user selects the filter value ‘Calculated’ because only a subset of individuals have a polygenic risk score calculated.

On the hereditary cardiovascular conditions dashboard (https://data.color.com/v2/cardio.html), users can apply the same types of query filters as those available on the hereditary cancer dashboard, with the following substitutions for clinical risk models and polygenic risk scores: ‘CHARGE - AF Risk Score’, ‘Framingham Risk Score’, ‘AF Polygenic Risk Score’, and ‘CAD Polygenic Risk Score’. For ‘AF Polygenic Risk Score’, and ‘CAD Polygenic Risk Score’, results will only be displayed if a user selects the filter value ‘Calculated’ because only a subset of individuals have polygenic risk scores calculated. These filter categories and filter values are listed in Table 2. The same ‘AND’ logic and ‘OR’ logic apply, as described above.

### Population characteristics

The population characteristics of the hereditary cancer dataset in Color Data v2 are very similar to those in Color Data v1. However, due to changes in inclusion criteria, there are some notable differences. These include an increase in the proportion of men (27.6% versus 20.4%) and non-Causasian individuals (30.6% versus 27.9%). The frequency of pathogenic variants in the total population increased to 11.0% in Color Data v2, compared to 10.8% in Color Data v1. There are a total of 14,269 unique variants in Color Data v2. The newly added query results show that 9.3% of individuals have an elevated, five-year risk of breast cancer as estimated using the Gail Model, and only 0.8% have an elevated lifetime risk of breast cancer as estimated using the Claus Model.

The 18,783 individuals in the new hereditary cardiovascular conditions dataset are a subset of the individuals in the hereditary cancer dataset. The majority of individuals are female (68.5%) and reported Caucasian ethnicity (70.4%). The average age at the time of genetic testing was 46.2 years. A total of 14,213 (75.9%) individuals reported no personal history of cardiovascular disease and/or events. Approximately 190 (1.0%) individuals reported a personal history of having a stroke, and 170 (0.9%) individuals reported having a heart attack. A total of 2,464,482 variants were identified in 30 genes associated with hereditary cardiovascular conditions, with the largest percentages in *RYR2, MYH11*, and *DSP*. There are 9,727 unique variants, over half of which are benign or likely benign (54.4%). The frequency of pathogenic and likely pathogenic variants in the total population is 1.4%. Finally, 0.3% of individuals are categorized as being at high-risk for atrial fibrillation using the CHARGE-AF simple score, and 7.5% are estimated to have a high 10-year risk for coronary heart disease using the simple office-based Framingham Coronary Heart Disease Risk Score.

### Sample query 1: frequency of pathogenic variants in genes associated with hereditary cardiovascular conditions

Cardiovascular disease is a leading cause of death in the United States, accounting for one-third of deaths worldwide (15). Many individuals with hereditary cardiovascular conditions progress asymptomatically, and as a result, go undiagnosed until they present with a sudden cardiac event. Users can investigate the frequency of pathogenic variants in genes associated with hereditary cardiovascular conditions in the database by filtering ‘Classification: Pathogenic or Likely pathogenic’ (https://data.color.com/v2/cardio.html#classification=Likely%20pathogenic&classification=Pathogenic). A total of 263 individuals have a pathogenic or likely pathogenic variant, the majority of which are female (64.6%) (Figure 1A) and Caucasian (64.6%) (Figure 1B). The average age at testing was 44.2 years (Figure 1A), and the majority of individuals reported no personal history of cardiovascular disease and/or events (67.7%, n=178) (Figure 1C). Nearly one-third of variants were identified in *MYBPC3* (30.5%), followed by *LDLR* (19.6%), *KCNQ1* (8.3%), and *PKP2* (8.3%) (Figure 1D). Of the 266 pathogenic or likely pathogenic variants identified, the most common variant was *MYBPC3* c.3628-41_3628-17del (n=38) (Figure 1E), which is associated with hypertrophic cardiomyopathies and found at high frequencies (from 2-8%) in Indian populations (16). To investigate the subpopulation of individuals with this variant, users can filter by ‘Gene: *MYBPC3*’ and ‘Variant: c.3628-41_3628-17del’ (https://data.color.com/v2/cardio.html#gene=MYBPC3&variant=c.3628-41_3628-17del). In this subpopulation of 40 individuals, 85.0% are of Indian descent (Figure 1F). Of those who provided information about their personal history (n=29), no one reported having a personal history of cardiovascular disease and/or events (Figure 1G). Taken together, researchers could use this data to better characterize the prevalence of hereditary cardiovascular disorders in a younger, unaffected population to identify asymptomatic individuals who are at risk for future cardiovascular disease and/or events.

**Figure 1.**
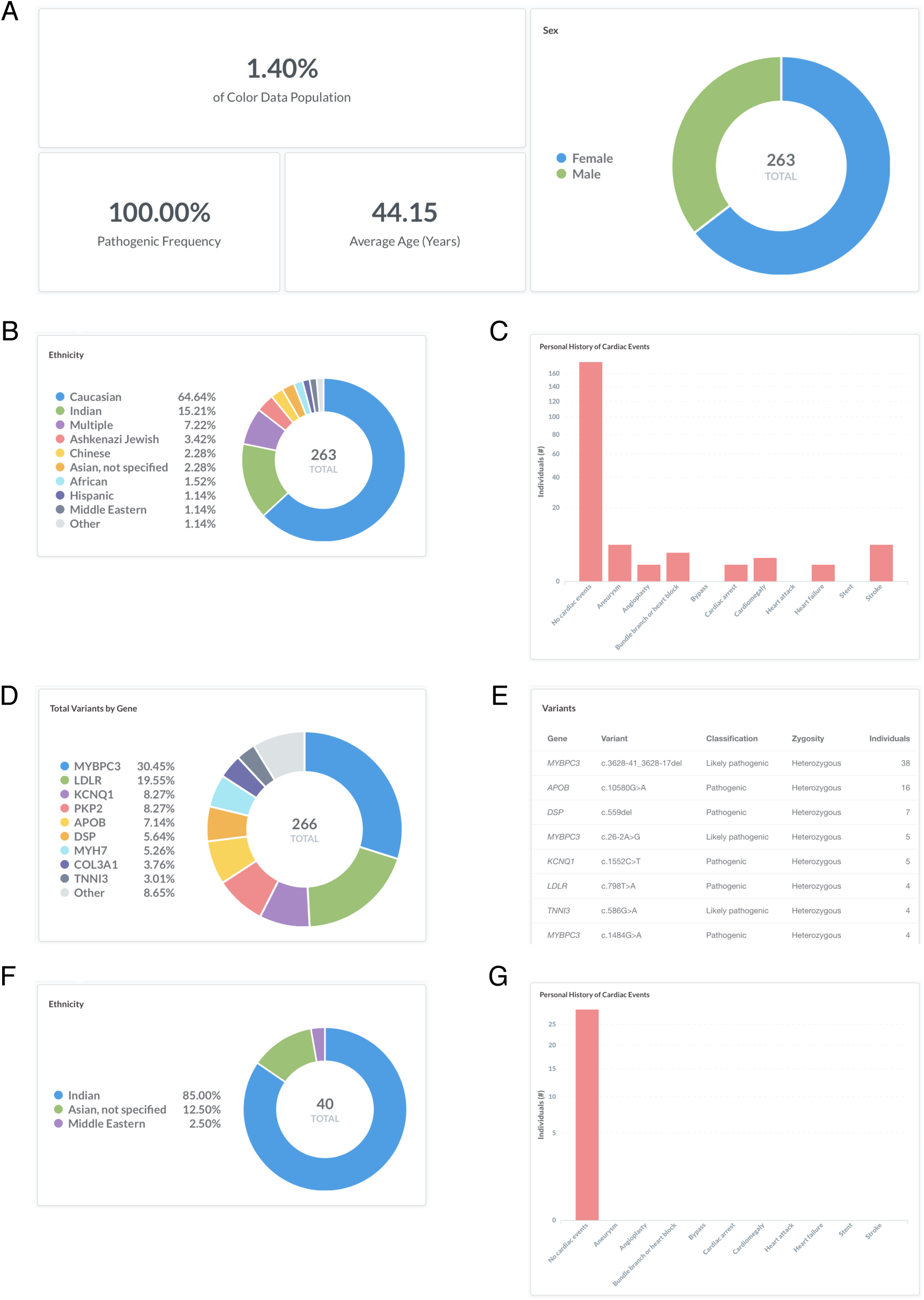
Screenshots of query results for frequency of pathogenic variants in genes associated with hereditary cardiovascular conditions. (A - E) Filter by ‘Classification: Pathogenic or Likely pathogenic’. Query URL: https://data.color.com/v2/cardio.html#classification=Likely%20pathogenic&classification=Pathogenic (F, G) Remove ‘Classification: Pathogenic or Likely pathogenic’ and filter by ‘Gene: *MYBPC3*’ and ‘Variant: c.3628-41_3628-17del’. Query URL: https://data.color.com/v2/cardio.html#gene=MYBPC3&variant=c.3628-41_3628-17del

### Sample query 2: monogenic and polygenic breast cancer risk in women with a personal history of breast cancer

Breast cancer is a common, complex disease that is associated with rare pathogenic and likely pathogenic variants (‘monogenic risk’) and the cumulative effect of many common changes across the genome (‘polygenic risk’) (9,10). Recent work has suggested that monogenic risk and polygenic risk interact to modify an individual’s overall risk for disease (17–19). To investigate monogenic and polygenic risk in women with a personal history of breast cancer, users can filter by ‘Sex: Female’, ‘Personal health history: Breast’, and ‘BC Polygenic Risk Score: Calculated’ (https://data.color.com/v2/cancer.html#sex=Female&personal_health_history=Breast&bc_polygenic_risk_score=Calculated). A total of 1443 females in the dataset reported a personal history of breast cancer and had a polygenic risk score for breast cancer risk calculated (Figure 2A).

**Figure 2.**
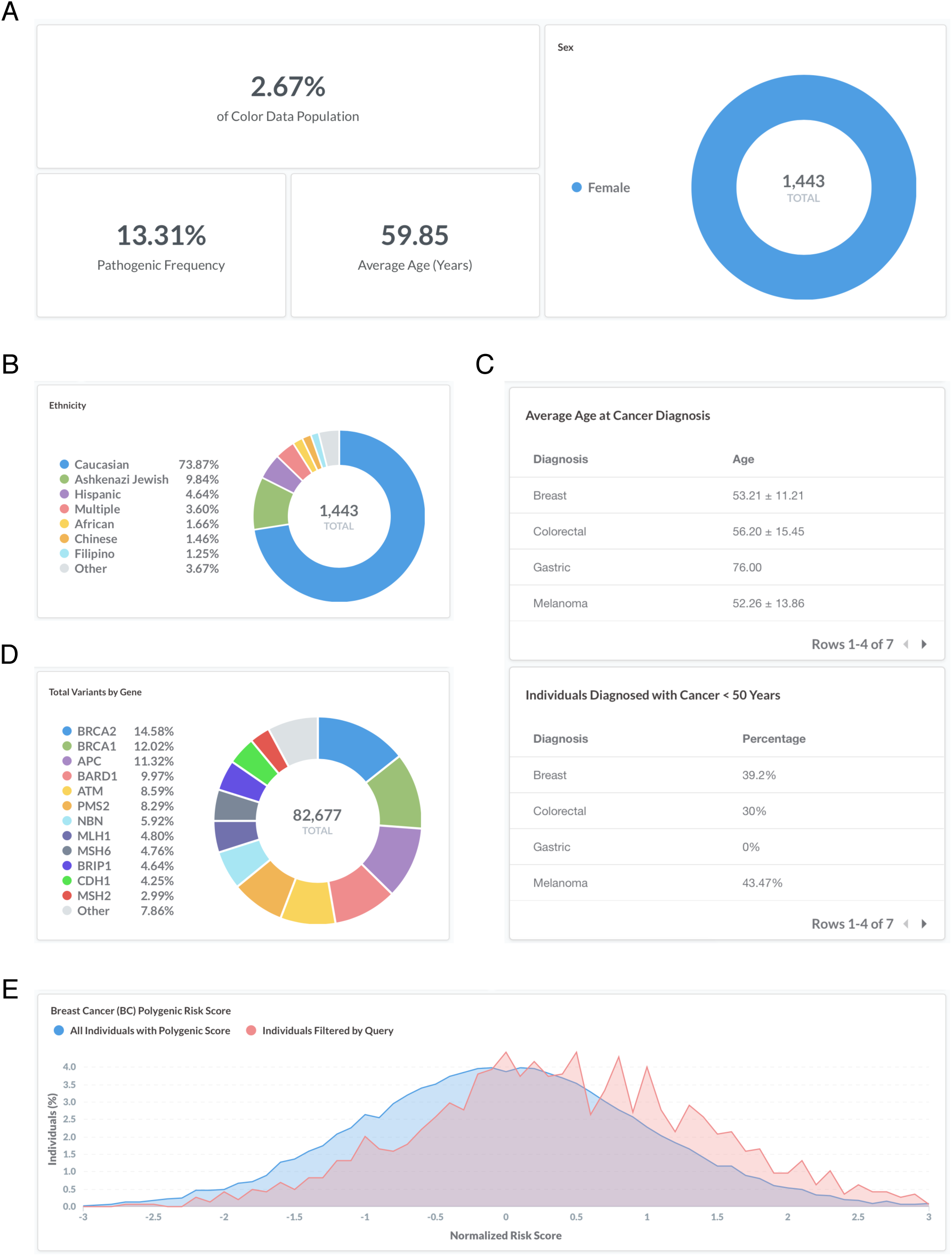
Screenshots of query results for monogenic and polygenic breast cancer risk in women with a personal history of breast cancer. (A - E) Filter by ‘Sex: Female’, ‘Personal health history: Breast’, and ‘BC Polygenic Risk Score: Calculated’. Query URL: https://data.color.com/v2/cancer.html#sex=Female&personal_health_history=Breast&bc_polygenic_risk_score=Calculated

The majority of individuals are Caucasian (73.9%) (Figure 2B), with an average age of 59.9 years old at the time of genetic testing (Figure 2A). The average age of diagnosis for breast cancer was 53.2 ± 11.2 years (standard deviation), and 39.2% of females were less than 50 years old at the time of diagnosis (Figure 2C). The pathogenic frequency in this population was 13.3% (Figure 2A). A total of 82,677 variants were identified, with the majority of variants in *BRCA2* (14.6%), *BRCA1* (12.0%), and *ATM* (11.3%) (Figure 2D). Compared to the normal distribution of risk scores among all individuals in the dataset (in blue), the distribution in women with a personal history of breast cancer is left-skewed (in red) (Figure 2E). Taken together, users could use this data to investigate the risk conferred by monogenic and polygenic risk factors in women with a history of breast cancer.

## DISCUSSION

In Color Data v2, we added new query filters and results such as family health history as well as clinical risk models and a polygenic score for breast cancer to the existing hereditary cancer dataset. Overall, the total number of individuals in this dataset increased to 54,000, however, the individuals in Color Data v1 are not a strict subset of the individuals in Color Data v2. This is due to a difference in inclusion criteria between the two versions. In Color Data v1, individuals were included in the database if they had received clinical genetic testing for all or any subset of the 30 hereditary cancer genes. In Color Data v2, individuals were only included if they received genetic testing for all 30 genes. This change in cohort composition likely contributed to the observed change in frequency of pathogenic and likely pathogenic variants in the total population and the increase in the number of total and unique variants.

Database users can also now explore a new disease area with self-reported phenotypic information and genetic data for 30 genes associated with hereditary cardiovascular conditions from 18,738 individuals. The frequency of pathogenic and likely pathogenic variants in our dataset was higher than previously reported estimates (20,21). This could be due to the generally younger age of individuals in the cohort and/or reduced penetrance in asymptomatic carriers. Genetic testing for hereditary cardiovascular conditions at population-scale has only recently begun, and sharing results through genetic databases such as Color Data will help rapidly advance our understanding of cardiovascular disease risk. Coronary artery disease may be of particular interest given the influence lifestyle modifications have been shown to have on lowering risk for disease. In a prospective study of more than 55,000 individuals, it was found that a healthy lifestyle was associated with significantly reduced risk of cardiovascular events across all genetic risk groups (22).

Similar to Color Data v1, Color Data v2 may be limited by selection bias for Caucasians and women, as well as by self-reported phenotypic information. Not all individuals in the database provided enough information to calculate risk for the newly included clinical risk models or had polygenic risk scores calculated, resulting in incomplete datasets. As the field continues to evaluate the personal and clinical utility of polygenic risk scores, it will be important to consider their predictive power in light of other risk factors. In addition, the clinical risk models and polygenic scores shown may change over time as more evidence emerges and novel models are discovered.

## DATA SHARING STATEMENT

The data in this report are publicly available at Color Data (https://data.color.com/). All reported variants have been submitted to ClinVar (https://www.ncbi.nlm.nih.gov/clinvar/submitters/505849/).

## ACKNOWLEDGEMENTS

We would like to thank Carman Lai and Sydney Okumura for insightful user-testing and Michael

K. Doney, Julian R. Homburger, Lauren Ryan, and Stephanie E. Wallace for helpful discussions.

